# Measuring amber initiator tRNA orthogonality in a genomically recoded organism

**DOI:** 10.1101/524249

**Authors:** Russel M. Vincent, Bradley W. Wright, Paul R. Jaschke

**Author notes:** Correspondence and requests for materials should be addressed to PRJ.

## Abstract

Using engineered initiator tRNA for precise control of protein translation within cells has great promise within future orthogonal translation systems to decouple housekeeping protein metabolism from that of engineered genetic systems. Previously, *E. coli* strain C321.ΔA.*exp* lacking all UAG stop codons was created, freeing this ‘amber’ stop codon for other purposes. An engineered ‘amber initiator’ 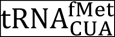 that activates translation at UAG codons is available, but little is known about this tRNA’s orthogonality. Here, we combine for the first time the amber initiator 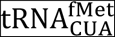in C321.ΔA.*exp* and measure its cellular effects. We found that the 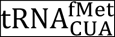expression resulted in a nearly 200Yfold increase in fluorescent reporter expression with a unimodal population distribution and no apparent cellular fitness defects. Proteomic analysis revealed upregulated ribosomeYassociated, tRNA degradation, and amino acid biosynthetic proteins, with no evidence for offYtarget translation initiation. In contrast to previous work, we show that UAGYinitiated proteins carry NYterminal methionine exclusively. Together, our results identify beneficial features of using the amber initiator 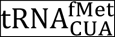to control gene expression while also revealing fundamental challenges to using engineered initiator tRNAs as the basis for orthogonal translation initiation systems.

---

The ability to precisely control an orthogonal protein translation system within engineered organisms is one of the grand goals of synthetic biology.^1^ A distinct feature of the standard genetic code is the dual function of AUG as both the dominant start codon as well as a methionine sense codon internal to genes. This dual function of the AUG start codon can create ambiguities in gene prediction and genetic design. For example, in recombinant protein design, inYframe AUG sense codons can behave as start codons if the RNA sequence upstream resembles a ShineYDalgarno site.^2^ Therefore, engineering an orthogonal system to initiate translation at a unique codon reserved solely for that function would be beneficial. Once established, the unique start codon could be paired with an orthogonal initiator tRNA aminoacylated with natural or unnatural amino acids with useful properties as has been done before in internal codons.^3^

Specific tRNAs recognise and decipher mRNA codons during translation and sequentially associate a bound amino acid to a growing polypeptide within the ribosome through three anticodon nucleotides (positions 34Y36) on the anticodon loop. tRNAs are divided into two functionally and structurally distinct classes: initiator tRNAs and elongator tRNAs. Initiator tRNAs participate in the initiation phase of translation, recognizing start codons, and thus, incorporate the first amino acid. The three consecutive G:C base pairs in the tRNA anticodon stem is a crucial conserved element for initiator tRNA binding to the PYsite in the 30S ribosomal subunit via interactions with the 16S ribosomal RNA.^4Y5^ Additionally, the initiator tRNA’s conserved C1:A72 base pair mismatch in the acceptor stem specifically interacts with methionylYtRNA NY formyltransferase enzyme^6^, and initiation factor 2 (IF2)^7^, a protein vital for prokaryotic translation initiation.

Over the past two decades, efforts have focused on engineering stop codon suppression systems that use elongator tRNA aminoacylYsynthetase pairs as an orthogonal system that specifically recognises amber (UAG) sense codons to siteY specifically incorporate unnatural amino acids ^8^. Unnatural amino acid incorporation enables researchers to build engineered proteins with an extended range of structural and functional properties for applications in protein evolution^9^, therapeutics^10^, and elucidating structure and function^11^. To prevent undesired natural amber codon suppression, competition for UAG codons from release factor 1, and nonYspecific unnatural amino acid incorporation, researchers have recently built a genomically recoded *Escherichia coli* strain, C321.ΔA with all instances of UAG stop codons and associated release factor (RF1) removed.^12^

The UAG amber stop codon has also been repurposed as a unique initiation codon in *E. coli* by creating an initiator 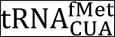with anticodon changed from the canonical CAU to CUA. This amber initiator has been shown in chloramphenicol acetyltransferase (CAT) reporter assays on bulk cell cultures to initiate translation *in vivo* from a UAG start codon.^13^ They found that initiation efficiency was only 50Y60% relative to the AUG reporter and that it was aminoacylated predominantly by glutamine instead of methionine.^13Y14^ Despite excellent prior work characterizing the amber initiator tRNA, a more complete picture of its interactions with the host cell is needed if an orthogonal translation system is to be based on this tRNA.

In this work, we characterized the phenotypic effects of expressing the amber initiator 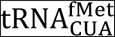in the genomically recoded C321.ΔA.*exp* strain using a modular tRNA/reporter plasmid system. Using this system we found that expressing 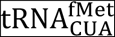in *E. coli* C321.ΔA.*exp* resulted in initiation from a UAG start codon with similar populationYlevel characteristics to that seen from the canonical AUG initiation codon and wildYtype initiator tRNA. We also found previously uncharacterised effects on host proteome, but surprisingly, found no evidence for offYtarget translation initiation from genomic UAG codons. Additionally, we found that proteins initiated from 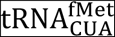carried methionine exclusively at their NYterminus. Together, this work reveals specific opportunities and challenges to using engineered initiator tRNAs in orthogonal translation systems.

## RESULTS AND DISCUSSION

To measure orthogonality of the previously described amber initiator 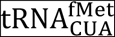 we built a modular plasmidYbased amber initiator tRNA expression and reporter system (Figure 1A). The modular system consists of the amber initiator plasmid (pULTRA::*tac2 metY*(CUA)) (Figure 1A, top) harbouring the *metY*(CUA) gene expressing 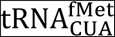^13^ and a compatible reporter plasmid pQEY60::T5Y*sfGFP* (Figure 1A, bottom) encoding one of three superfolder green fluorescent protein (sfGFP) variants differing in their start codon (either AUG, UAG, or GCC), where sfGFP(AUG) denotes the superfolder GFP gene with AUG start codon. To reduce host interference from release factor 1 competing with UAG codons we chose *E. coli* C321.ΔA.*exp* (Table 1) with a reduced genetic code as the host strain for this work.^12^

**Table 1.** Description of strains and plasmids used in this study.

| Strain or Plasmid | Description | Reference or Source |
| --- | --- | --- |
| Strain |  |  |
| C321.ΔA.exp | *E. coli* MG1655 Δ(*ybhB-bioAB*)::*zeoR* Δ*prfA*; all 321 UAG stop codons changed to UAA. | Addgene #49018 |
| Plasmids |  |  |
| pULTRA::*tac-metY*(CUA) | pULTRA plasmid with *metY* gene containing amber initiator anticodon (CUA) driven by *tacI* promoter. | 35 and this study |
| pULTRA::*tac-metY*(CAU) | pULTRA plasmid with *metY* gene containing wild-type initiator anticodon (CAU) driven by *tacI* promoter. | 35 and this study |
| pULTRA::*tac*-Empty | pULTRA plasmid with *tacI* promoter, lacking any insert. | 35 and this study |
| pQE-60::T5-*sfGFP*(UAG) | pQE-60 plasmid with *sfGFP* gene initiated by UAG start codon and C-terminal 6xHis-tag inserted into the plasmid's multiple cloning site. | Qiagen and this study |
| pQE-60::T5-*sfGFP*(AUG) | pQE-60 plasmid with *sfGFP* gene initiated by AUG start codon and C-terminal 6xHis-tag inserted into the plasmid's multiple cloning site. | Qiagen and this study |
| pQE-60::T5-*sfGFP*(GCC) | pQE-60 plasmid with *sfGFP* gene initiated by GCC start codon and C-terminal 6xHis-tag inserted into the plasmid's multiple cloning site. | Qiagen and this study |
| pSMART-BAC::*pRha-NanoLuc*(UAG) | pSMART-BAC plasmid with single-copy origin of replication (*oriS*, *repE*, *parABC*); TrfA-inducible high-copy origin of replication (*oriV*); Cm<sup>r</sup>-chloramphenicol resistance gene; NanoLuc gene with UAG start codon under control of rhamnose promoter (*pRha*) inserted into the vector's multiple-cloning site. | Lucigen Corp. and this study |
| pSMART-BAC::*pRha-NanoLuc*(AUG) | pSMART-BAC plasmid with single-copy origin of replication (*oriS*, *repE*, *parABC*); TrfA-inducible high-copy origin of replication (*oriV*); Cm<sup>r</sup>-chloramphenicol resistance gene; NanoLuc gene with AUG start codon under control of rhamnose promoter (*pRha*) inserted into the vector's multiple-cloning site. | Lucigen Corp. and this study |

**Figure 1.**
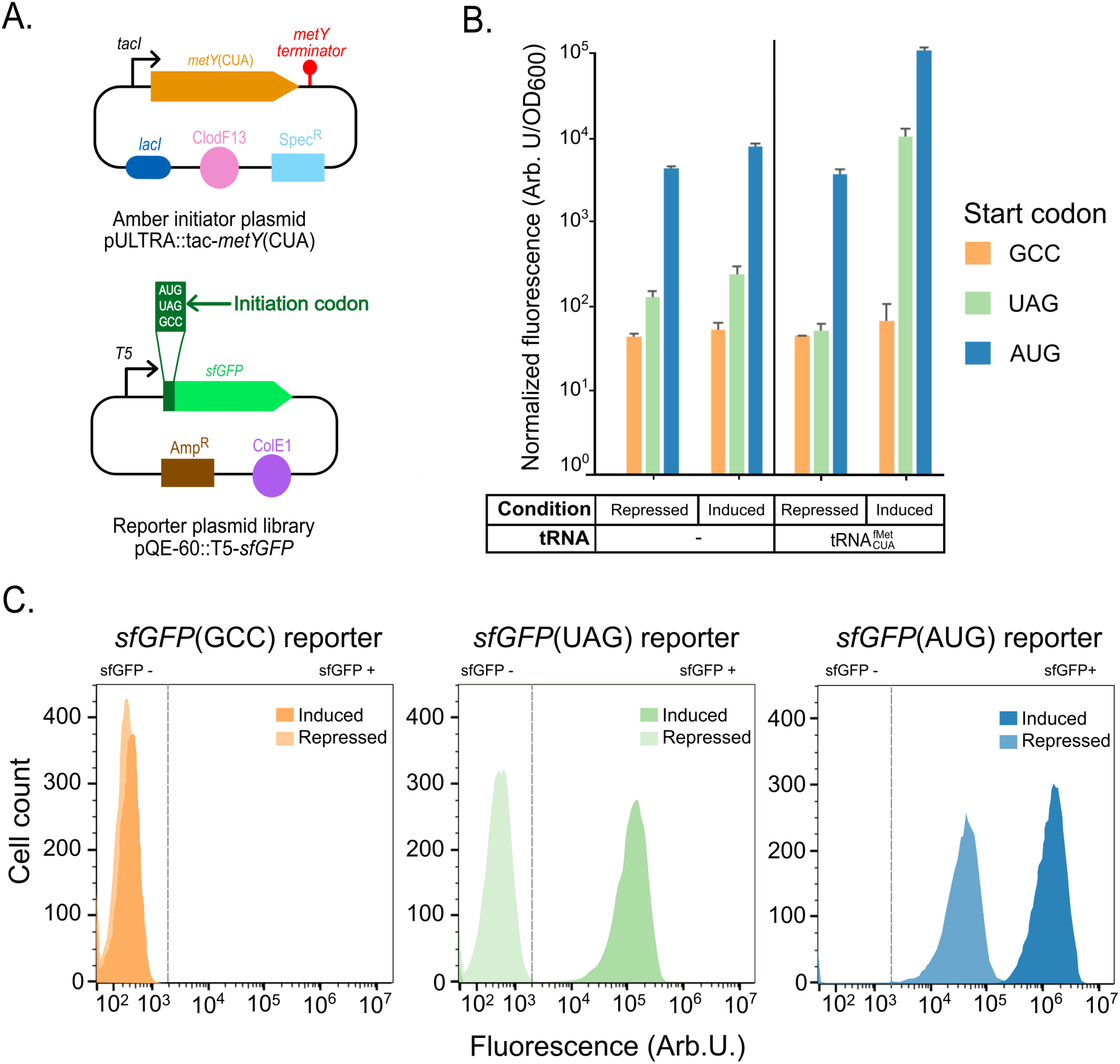
Expression of amber initiator tRNA causes increase in reporter fluorescence from both amber UAG and canonical AUG start codons. (A) Amber initiator and reporter plasmid system design. Amber initiator plasmid pULTRA*::tac2metY*(CUA) transcribing 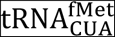, and reporter plasmid pQEY60 library expressing sfGFP containing three different initiation codons (AUG, UAG, and GCC). (B) Normalized expression levels from sfGFP reporters under different 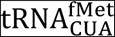transcription conditions. Repressed, 2% glucose; Induced, 1 mM IPTG; tRNA, presence or absence of 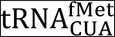expressing amber initiator plasmid. Average of three biological replicates with error bars showing one standard deviation. (C) Representative fluorescence (Arb. U.) histogram of gated populations containing pULTRA*::tac2metY*(CUA) expressing 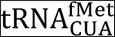with pQEY60 reporter library. Induced cell population (darker shades) and repressed cell population (lighter shades) shown for each reporter. Vertical dashed lines demarcate fluorescent and nonYfluorescent populations based on the 99^th^ percentile of the control population (fluorescence of *E. coli* C321.∆A*.exp* population expressing 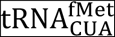with sfGFP(GCC) reporter).

We used this plasmid system to measure translation initiation events due to 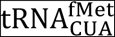expression. In the absence of pULTRA::*tac2metY*(CUA) plasmid, the sfGFP(UAG) reporter fluorescence was indistinguishable from the negative control sfGFP(GCC) fluorescence (Figure 1B). However, when the 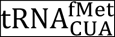was present in the cell, sfGFP(UAG) reporter fluorescence increased 196Yfold in the induced condition compared to the repressed condition (Figure 1B and Figure S1). Moreover, 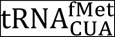expression resulted in sfGFP(UAG) fluorescence at a similar level to that of sfGFP(AUG) fluorescence when 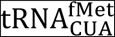was absent in the system (Figure 1B and Table S1). Fluorescence from reporter sfGFP(GCC) remained nearly unchanged in all conditions, consistent with previous findings.^15^ These data confirm the ability of 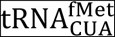to efficiently initiate protein synthesis from the UAG stop codon at nearY cognate AUG levels.^13^

Surprisingly, 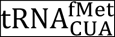expression caused an increase in the sfGFP(AUG) fluorescence per cell (Figure 1B, right blue bars) suggesting the possibility that 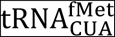could also recognise and interact with AUG start codons. Previous work concluded that the wildYtype host initiator tRNA (with a CAU as opposed to CUA anticodon) does not initiate translation from a UAG start codon.^15^ The reciprocal interaction of a CUA anticodon with an AUG start codon should also result in weak translation initiation.

We next used flow cytometry to measure the distribution of fluorescence within the population of *E. coli* C321.ΔA.*exp* cells harbouring the pULTRA::*tac*Y*metY*(CUA) plasmid. All cell populations had unimodally distributed fluorescence in both repressed and induced conditions and had uniform populationYwide increase in fluorescence in both the sfGFP(UAG) reporter and the sfGFP(AUG) reporter strains upon 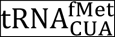induction (Figure 1C). This is the first measurement, to our knowledge, of populationY level translationYinitiation from the amber initiator tRNA.

We next performed growth assays to measure the fitness of cells harbouring pULTRA::*tac2metY*(CUA). We compared growth characteristics of cells harbouring plasmids expressing the amber initiator tRNA (pULTRA::*tac2metY*(CUA)) and the wildY type initiator tRNA (pULTRA::*tac2metY*) to the empty vector (pULTRA::*tac*YEmpty) under inducing conditions. We found no significant effect (pYvalue > 0.5) from expression of either initiator tRNA on maximal growth rate or maximal cell density (Figure 2 and Table S2).

**Figure 2.**
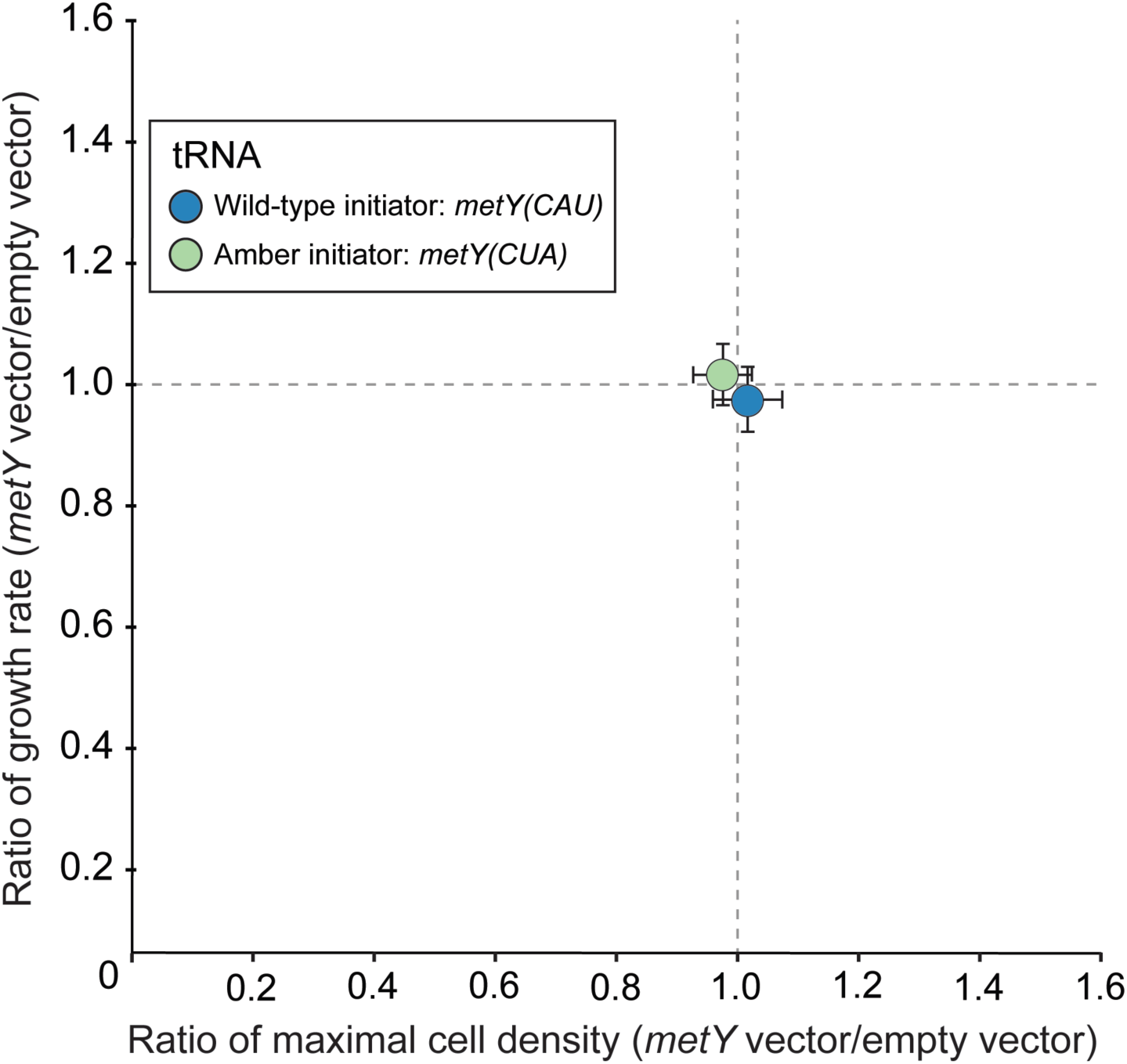
Amber initiator and wildYtype initiator tRNA expression results in minimal fitness impact on C321.ΔA*.exp* cells. Ratios of maximal cell densities (horizontal axis) and growth rate (vertical axis) were determined for strains containing either pULTRA*::tac2metY*(CUA) or pULTRA*::tac2metY*(CAU) plasmid versus their corresponding strains with the empty vector pULTRA*::tac2*Empty. Strains unaffected by initiator tRNA expression are expected to have ratios of 1. Strains exhibiting slower growth are below the horizontal gray line, and strains exhibiting lower maximum cell density are to the left of the vertical gray line. All strains were grown under the same induced (1 mM IPTG) condition. Each color represents the average of biological replicates (n = 3), and error bars show one standard deviation.

To better understand the global proteomic effect of expressing the amber initiator tRNA in C321.ΔA.*exp,* as well as to investigate possible causes for the observed increase in AUG reporter fluorescence (Figure 1), we next measured global protein production in these cells. We employed the dataYindependent acquisition method SWATHYMS (Sequential Window Acquisition of all Theoretical fragment ion Mass Spectra) which enables the collection of large, accurate, and reproducible quantitative proteomic datasets.^16Y17^ We collected data from *E. coli* C321.ΔA.*exp* samples with and without 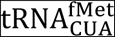expression three and five hours postYinduction.

We first interrogated the SWATHYMS data set with the aim of detecting canonical *E. coli* proteins with altered abundance under 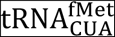expression. Summed fragment ion peak areas (area under the curve) of measured precursor ions (peptides) allowed us to compare protein abundance between *E. coli* C321.ΔA.*exp* expressing 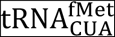versus the control. Three hours postYinduction we identified only two differentially expressed proteins that reached our significance threshold (foldYchange >1.5, pYvalue <0.01). The lactose operon repressor protein (LacI) was upYregulated 4.9Yfold because it was constitutively expressed from the mediumYcopy pULTRA::*tac*Y*metY*(CUA) plasmid (Figure 3A, brown point and Table S3). The FhuE receptor protein was also upY regulated 3.7Yfold (Figure 3A, purple point). Together, these data show that at three hours postYinduction, the amber initiator 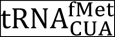does not appear to have a largeYscale effect on the C321.ΔA.*exp* proteome.

**Figure 3.**
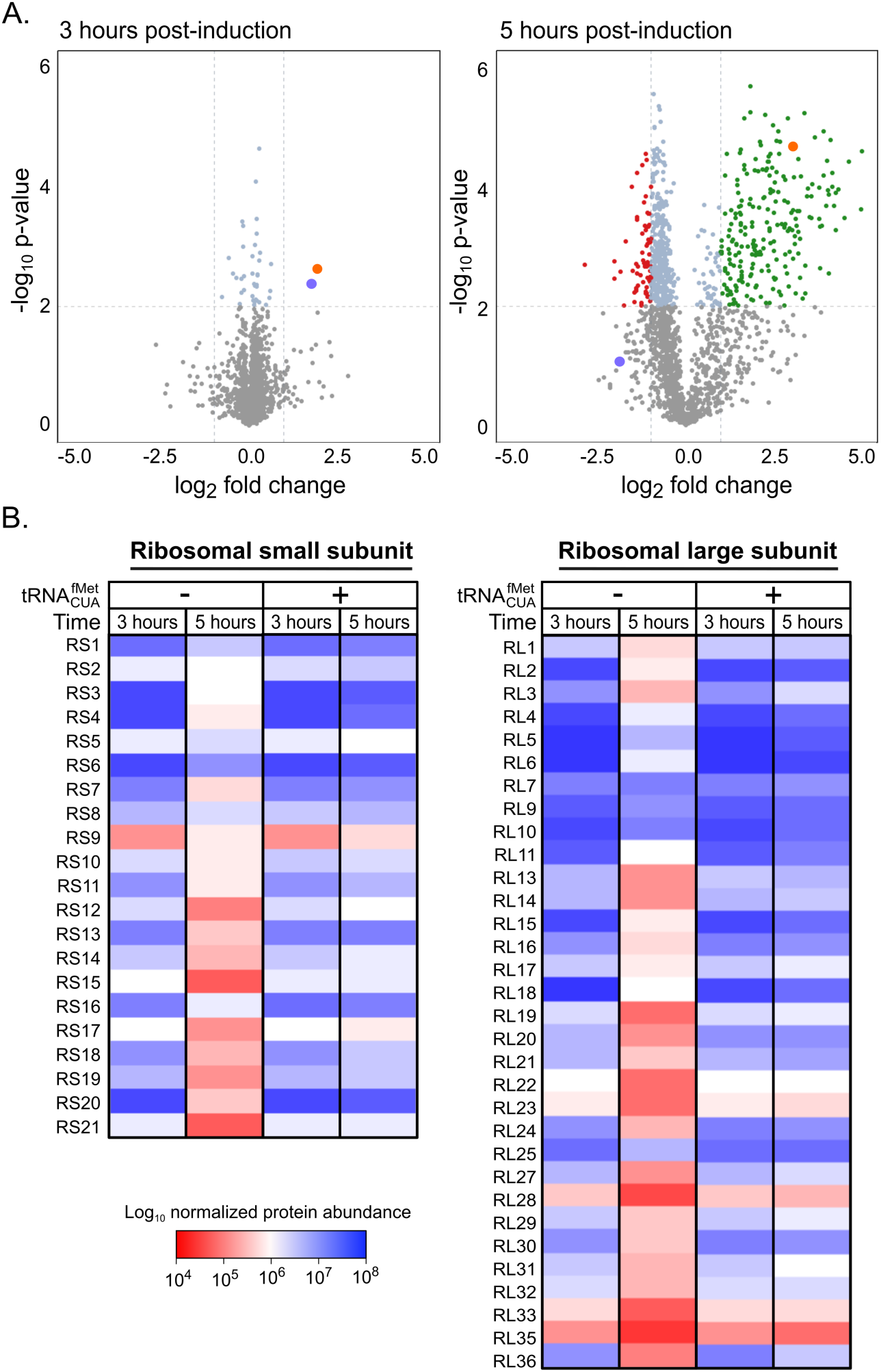
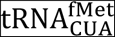expression causes delayed downregulation of ribosomal proteins in C321.ΔA*.exp* cells. (A) Volcano plot of 1,803 quantified proteins from *E. coli* C321.ΔA*.exp* cells at 3Yhours and 5Yhours after amber initiator 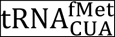induction. The orange points represent LacI and purple points represent FhuE. Vertical grey dashed lines represent fold change threshold: log_2_ ≥ 1 or ≤ −1 while the horizontal grey dashed line represents significance threshold: Y log_10_ pYvalue ≥ 2 equivalent to pYvalue ≤ 0.01. Green points represent significantly upYregulated proteins, red point represent significantly downY regulated, blue points represent unaltered expression, and grey points represent proteins that did not reach the significance threshold (pYvalue ≥ 0.01). (B) HeatYmap of ribosomal proteins quantified in C321.ΔA*.exp* control cells versus C321.ΔA*.exp*(pULTRA::*tac*Y*metY*(CUA)) cells expressing 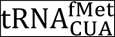. The C321.ΔA*.exp* control and C321.ΔA*.exp*(pULTRA::*tac*Y*metY*(CUA)) strains were grown in LB in the presence of 1 mM IPTG and harvested at three and five hours postYinduction.

In contrast to three hours postYinduction, at five hours we identified 262 differentially expressed proteins (Figure 3A and Table S4). To assess the biological relevance of the differentially expressed proteins, we performed enrichment analysis employing the Protein ANalysis THrough Evolutionary Relationships (PANTHER) functional annotation tool.^18^ This analysis revealed that 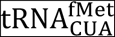expression had a significant impact on levels of proteins involved in the ribosome and translation process, leading to enriched biological process categories such as ribosomal biogenesis and assembly, and peptide metabolic pathways (Figure S2 and Table S5). Our data also showed that ribosomal proteins were abundant in both 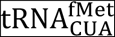-expressing and control cells three hours postYinduction, but that they declined five hours postY induction in control cells. In contrast, when 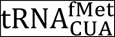was expressed in the cell, ribosomal protein levels did not decline (Figure 3B and Figure S3, cluster 3).

In addition to the sustained production of the majority of ribosomal structural proteins five hours postYinduction, eight ribosomal RNA (rRNA) modification enzymes, two initiation factors (IF2 and IF3), and elongation factor 4 (LepA) were all significantly upYregulated 2.0 Y 9.1Yfold compared to control cells (Table S4). This expression profile was similar to ribosomal proteins that continued to be expressed at levels similar to three hours postYinduction while levels in control cells declined (Figure S3, cluster 3). Notably, LepA plays a direct role in ribosomal biogenesis^19^ and a simultaneous increase in observed rRNA modification proteins could suggest a rise in mature ribosome concentration. A recent study has linked ribosomal rRNA biogenesis to the conserved anticodon stem sequence of the initiator tRNA.^20^ Our results complement the findings of this previous study by showing that ribosomal protein is also highly upregulated in response to increased cellular levels of initiator tRNA. We speculate that an increased number of mature ribosomes, in addition to upYregulated initiation factors (IF2 and IF3), could explain the increased translation from the AUG start codon reporter we observed (Figure 1B, right blue bars), rather than 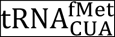directly interacting with AUG start codons.

Previous evidence showed that 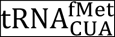is a poor substrate for aminoacylYtRNA synthetases^13Y14, 21^ and increases the level of uncharged tRNA in the cell^21^ triggering the tRNA catabolic pathways (Figure 4). Additionally, the levels of uncharged 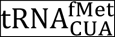may arise due to peptidyl tRNA hydrolase activity on amber initiator tRNA charged with glutamine.^22^ This potential imbalance in charged to uncharged tRNA ratio can be seen in our experiment through the increased abundance of poly(A) polymerase (PAP I) and polynucleotide phosphorylase (PNPase), both involved in tRNA degradation.^23^

**Figure 4.**
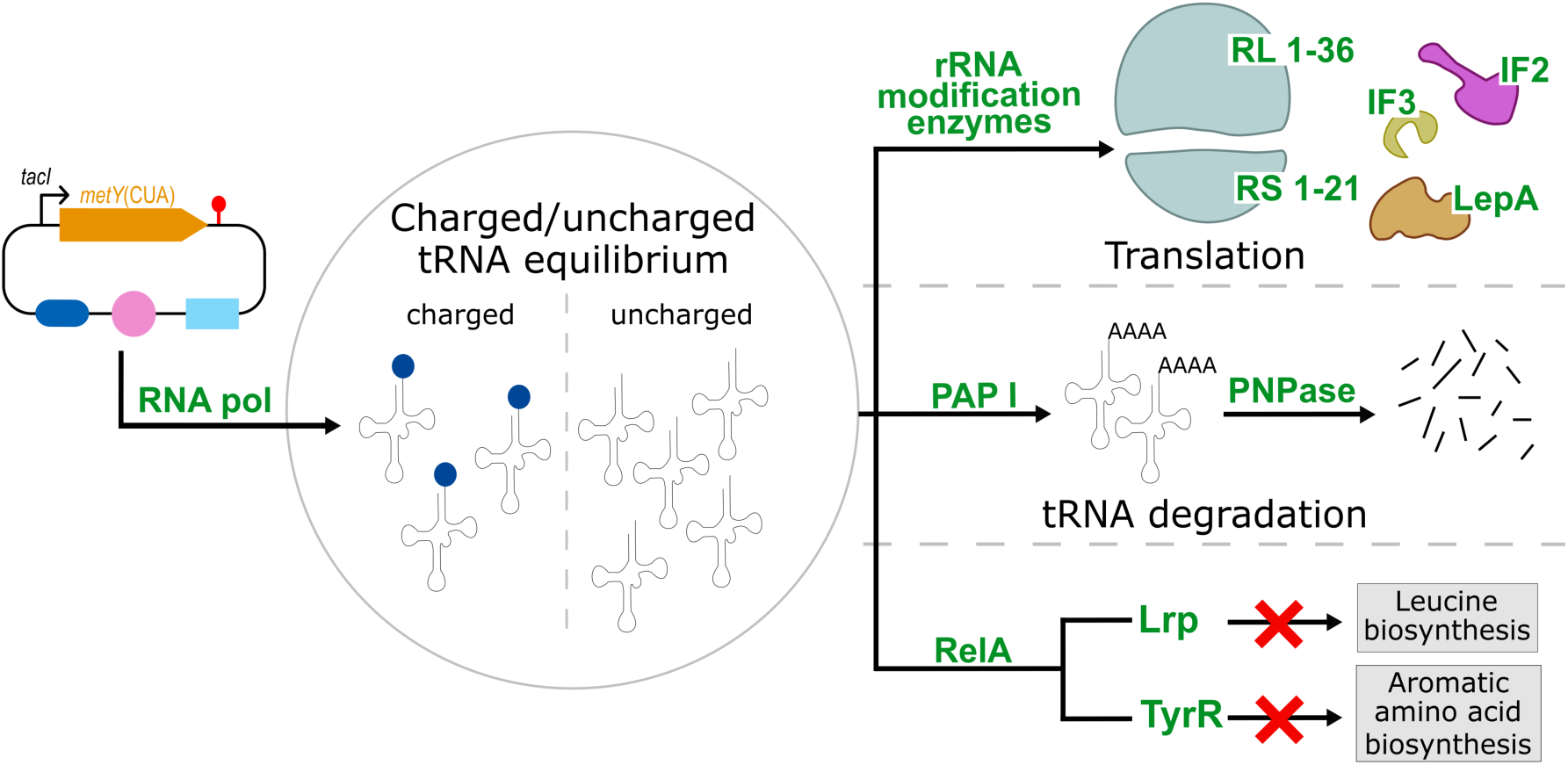
Proposed model of cellular effects of amber initiator 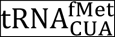expression. Increased RNA polymerase subunits transcribe 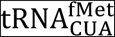from pULTRA::*tac*Y*metY*(CUA) resulting in reduction of charged 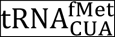. Black arrows represent differentially regulated pathways, green text represent upYregulated proteins, and gray boxes depict unaltered pathways.

We also observe an upregulation of guanosine pyrophosphatase (RelA)^24^ that is a key player in the cellular response to amino acid starvation through the stringent response.^25^ Despite RelA upregulation, it is unlikely that cells expressing the amber initiator tRNA are activating the stringent response, which is normally characterized by a reduction in the number of ribosomal proteins and expression of starvation sigma factors.^26^ Neither of these responses are observed in our experiment, and instead we see sigma factor 70 (σ^70^) increased 2.9Yfold and increased ribosomal protein expression (Table S4). We do observe the upYregulation of leucineYresponsive regulatory protein (Lrp) and transcriptional regulator TyrR, normally induced during the stringent response.^25^ Both Lrp and TyrR, in the absence of leucine and aromatic amino acids respectively, induce amino acid biosynthesis.^27Y28^ Providing further evidence that the stringent response is not being activated in response to the amber initiator tRNA expression, the increased levels of transcriptional regulators Lrp and TyrR in our work did not result in upYregulation of associated amino acid biosynthesis pathways normally observed in the stringent response.

We next searched our SWATH dataset for offYtarget peptides produced from open reading frames (ORFs) beginning with UAG start codons in the C321.ΔA*.exp* genome. The C321.ΔA.*exp* strain has 321 instances of functional UAG stop codons in the genome replaced with UAA stop codons^12^ but we hypothesized that of the 27,108 UAG codons remaining in the genome, some may initiate translation in the presence of the amber initiator 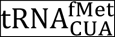. To identify peptides initiating from these UAG codons we generated an offYtarget proteome database using the genome sequence of C321.ΔA.*exp* (Figure 5A). We considered all genomic UAG codons as potential start codons and considered UGA and UAA as stop codons. We did not treat UAG as a stop codon because C321.ΔA.*exp* lacks release factor 1 which is required for termination at UAG stop codons. We generated a list of 27,108 ORFs and filtered out all ORFs shorter than 150 nucleotides (50 amino acids) in length since small proteins (less than 50 amino acids) are generally ignored in wholeYproteome studies to reduce background noise.^29^ This filtering process resulted in an offYtarget database of 9,857 ORFs initiating from UAG start codons. To further filter the ORF list we identified those ORFs with a ShineY Dalgarno (SD) sequence upstream of the UAG start codon, resulting in 990 ORFs in the final offYtarget database (Figure 5A and Table S6). Additionally, we considered that 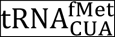could be charged with glutamine in addition to methionine^13Y14, 21^ and thus created two versions of each protein in the database, one with an NYterminal methionine and one with a glutamine.

**Figure 5.**
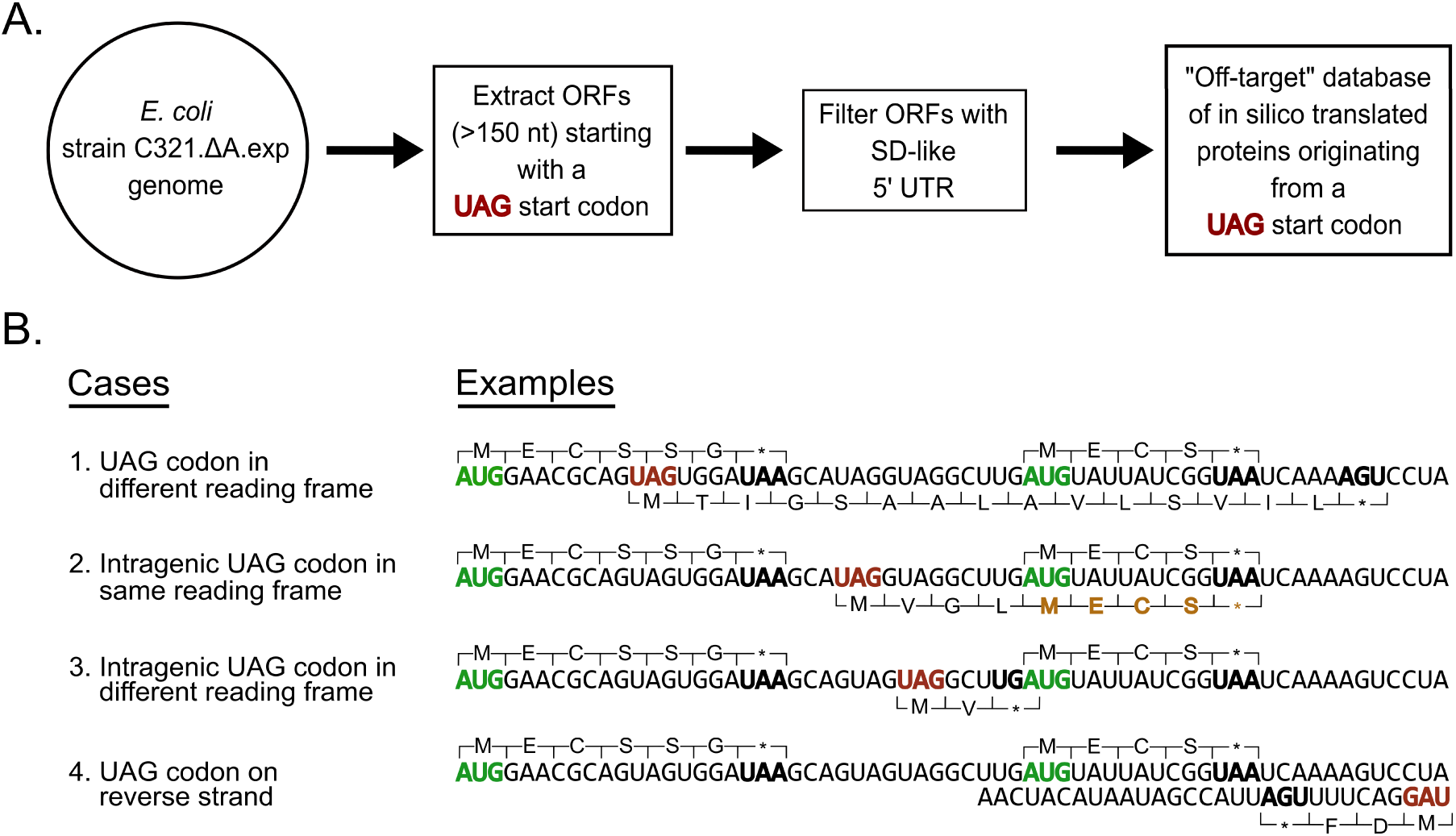
Method used to create UAG open reading frame offYtarget protein database. (A) Algorithmic workflow to create an offYtarget UAGYinitiating open reading frame (ORF) database from the C321.∆A*.exp* genome sequence. ORFs less than 150 nucleotides long were filtered out resulting in a reduction from 27,108 ORFs to 9,857. These ORFs were further filtered down to include only ORFs with ShineYDalgarnoYlike 5’ upstream sequences. The resulting 990 ORFs were then translated in two ways with UAG coding for methionine and glutamine to generate an offYtarget protein database. (B) UAG codon contexts that could result in ORFs with UAG start codon. The top proteins represent annotated sequences translated from canonical AUG start codon and bottom proteins represent hypothetical offYtarget protein sequences translated from a UAG start codon. ORFs displayed are for illustrative purposes only and are intentionally shorter than the minimum 150 nucleotide filter used to generate the offYtarget protein database. Green codons represent a canonical start codon; bold black codons represent a canonical stop codon in *E. coli* C321.∆A*.exp*; red codons represent a UAG start codon; brown amino acids represent shared peptides with sequences translated from a canonical AUG start codon and an amber UAG start codon.

We then combined the offYtarget database with the conventional *E. coli* MG1655 database (4,443 proteins) and searched our generated spectral library against it. Using the SWATHYMS data, we mined the combined spectral library and quantified 1,803 proteins across all three biological replicates that satisfied our significance criteria (protein and peptide level FDR <1% with at least 1 unique peptide observed). We observed 76 peptides that matched proteins from the offYtarget database, however, the identified peptides were shared with existing canonical *E. coli* proteins due to the presence of a UAG start codon upstream and inYframe with a canonical start codon (Figure 5B, case 2). We considered these peptides to originate from the conventional *E. coli* proteome since none were unique to the offYtarget database (Table S7). Despite not detecting any peptides originating from offYtarget ORFs we were able to detect peptides from four known proteins that have been previously shown to be present at 1Y5 copies per cell in *E. coli* grown in LB: P02942 (Tsr protein), P04949 (FliC protein), P07017 (Tar protein), and P39332 (YjgH protein).^30^ Together, these results show that despite detecting and quantifying over 1,800 proteins across a large dynamic range, we saw no evidence for 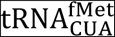expression leading to offYtarget translation.

To further probe the possibility of translation initiation from genomically encoded offYtarget ORFs with UAG start codons, we built a singleYcopy reporter plasmid to determine whether we could detect protein production at the low levels expected for these ORFs. We designed and built the bacterial artificial chromosome (BAC) plasmids pSMARTYBAC::*pRha*Y*NanoLuc*(UAG) and pSMARTYBAC::*pRha*Y*NanoLuc*(AUG) (Figure 6A) and transformed them it into cells already containing the pULTRA::*tac*Y*metY*(CUA) plasmid or with the empty vector pULTRA::*tac*YEmpty. We performed a luciferase assay on cells induced with varying concentrations of rhamnose designed to induce varying amounts of NanoLuc luciferase production with constant induction of the amber initiator 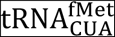or empty vector. As expected, we found that NanoLuc(UAG) luminescence was absolutely dependent on the amber initiator 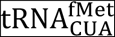expression (Figure 6B).

**Figure 6.**
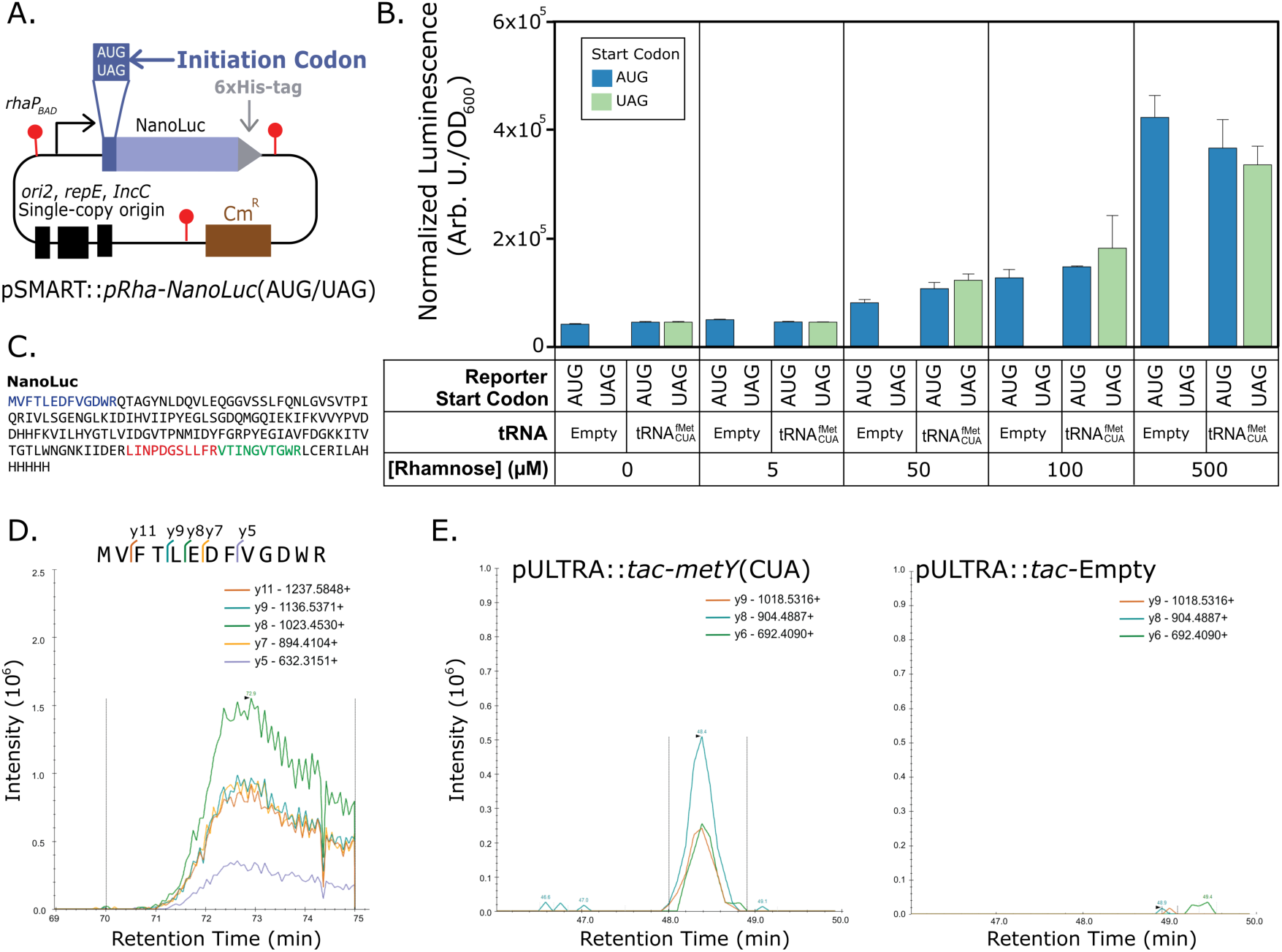
SingleYcopy bacterial artificial chromosome (BAC) reporter plasmid expressing NanoLuc shows NYterminal amino acid is methionine and that peptides are only observed when the amber initiator tRNA is coYexpressed with the reporter. (A) SingleYcopy BAC NanoLuc plasmid design. Red ball and sticks represent transcriptional terminators; *ori2*, *repE*, and *IncC* encode singleYcopy origin. (B) NanoLuc luciferase reporter expression from C321.ΔA.*exp* cells harbouring either pULTRA::*tac*YEmpty or pULTRA::*tac*Y*metY*(CUA) tRNA expression plasmids along with either pSMART::*pRha*Y *Nanoluc*(UAG) or pSMART::*pRha*Y*Nanoluc*(AUG) reporter plasmids. (C) Amino acid sequence of NanoLuc protein with peptides observed by PRM mass spectrometry highlighted in color. (D) Mass spectrometric identification of NYterminal amino acid of purified NanoLuc protein expressed with UAG start codon. Product ions specific to a methionyl NYterminal peptide from NanoLuc(UAG) shown with identical retention times and expected masses. Peptide sequence and preYcursor peptide fragmentation patterns shown above chromatogram. NanoLucY6xHis was NiYNTA purified from C321.ΔA.*exp*(pULTRA::*tac*Y*metY*(CUA); pSMART::*pRha*Y*NanoLuc*(UAG)) cells induced with 1 mM IPTG and 1 mM rhamnose. (E) NanoLuc peptides expressed from singleYcopy BAC detected in C321.ΔA.*exp* cells only when expressing the amber initiator. NanoLuc peptides observed in PRM mass spectrometry assay. Left, C321.ΔA.*exp*(pULTRA::*tac*Y *metY*(CUA); pSMART::*pRha*Y*NanoLuc*(UAG)) cells; Right, C321.ΔA.*exp*(pULTRA::*tac*Y Empty; pSMART::*pRha*Y*NanoLuc*(UAG)) cells.

At all rhamnose concentrations, when the amber initiator 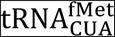was expressed, the luminescence resulting from NanoLuc(AUG) and NanoLuc(UAG) expression was equal (Figure 6B). We also found that the NanoLuc(AUG) reporter luminescence was independent of the presence of the amber initiator, having the same signal in the pULTRA::*tac*YEmpty cells as those containing pULTRA::*tac*Y*metY*(CUA).

This result is in contrast to what we found with the sfGFP reporter on a mediumY copy vector (Figure 1) where the presence of the amber initiator seemed to boost production of sfGFP(AUG). One explanation for this different behaviour could be that it is a result of the vector copy number: at lowYcopy number the mRNA encoding the NanoLuc protein is likely limiting production of the protein, whereas at much higher copy number the ribosomal abundance may be limiting sfGFP production. If this is true, and the expression of the amber initiator tRNA increases ribosomal abundance (Figure 3), then it follows that increased ribosome abundance in the medium copy sfGFP(AUG) reporter strain would cause an increase in fluorescence due to increased translation initiation from the reporter mRNA. In contrast, at lowYcopy increased ribosomal abundance has no effect on the NanoLuc(AUG) reporter because mRNA levels are limiting, not ribosomal abundance.

To identify the NYterminal amino acid incorporated into the NanoLuc reporter protein we purified the protein from C321.ΔA.*exp* cells harbouring the pSMARTY BAC::*pRha*Y*NanoLuc*(UAG) and pULTRA::*tac*Y*metY*(CUA) plasmids. We designed the pSMARTYBAC::*pRha*Y*NanoLuc*(UAG/AUG) plasmids to contain a CYterminal 6xHisYtag to enable purification of the fullYlength protein (Figure 6A). The internal and NYterminal peptides of the purified NanoLuc proteins were subjected to a targeted mass spectrometry technique called parallel reaction monitoring (PRM). The strength of the PRM method is that it provides high specificity and sensitivity and would allow us to specify peptides to target for accurate identification.^31^

The PRM assay showed that we could detect three peptides produced from tryptic digests of NanoLuc(UAG) (Figure 6C). The NanoLuc(UAG) NYterminal peptides contained methionine exclusively at the NYterminus (Figure 6D). The PRM assay was setup to agnostically target NYterminal peptides from NanoLuc(UAG) with any amino acid in the NYterminal position.^22^ This result is in contrast to previous work with the amber initiator tRNA, which showed both methionine and glutamine as the NYterminal amino acid from a reporter containing UAG as the start codon.^32^ However, these NY terminal amino acid differences could be a result of the different reporter gene used (NanoLuc versus CAT) or by the different bacterial strains used in the experiments (*E. coli* versus *Mycobacterium smegmatis)*.

Using the PRM method and the singleYcopy NanoLuc(UAG) reporter plasmid, we searched for offYtarget peptides in C321.ΔA.*exp* cells harbouring the pSMARTY BAC::*pRha*Y*NanoLuc*(UAG) and pULTRA::*tac*Y*metY*(CUA) plasmids. We also measured peptides from cells containing the pSMARTYBAC::*pRha*Y*NanoLuc*(UAG) and pULTRA::*tac2Empty* plasmids as a control. In addition to inducing the pULTRA plasmid, we also weakly induced the pSMARTYBAC::*pRha*Y*NanoLuc*(UAG) reporter with 100 µM rhamnose (Figure 6B) as an internal control to determine our ability to detect UAGY initiating proteins from singleYcopy genes. Since the PRM method derives its sensitivity from limiting the peptides the mass spectrometer measures to a preYspecified list, it was not possible to target all 990 UAGYinitiating ORFs with a ShineYDalgarno sequence in C321.ΔA.*exp*. Instead, we limited our search space to peptides from 194 offYtarget ORFs with the strongest upstream ShineYDalgarno sequences (Table S6), in addition to the peptides of NanoLuc(UAG). Simultaneously, we targeted four weakly expressed *E. coli* host proteins previously observed in our SWATHYMS dataset. As before, we found no evidence for offYtarget protein expression from any of the offYtarget ORFs. We did detect one peptide that could have originated from a UAGYinitiating ORF, but this peptide was observed in both the empty vector control and amber initiatorYcontaining cells (Figure S5) suggesting that expression of this peptide is not dependent on 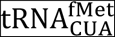. At the same time, we were able to detect peptides from the four weakly expressed proteins from the *E. coli* host proteome (Table S11) along with NanoLuc(UAG) reporter protein from the singleYcopy pSMART plasmid, but only when cells contained pULTRA::*tac*Y*metY*(CUA), not pULTRA::*tac*YEmpty (Figure 6E).

In conclusion, we found that the amber initiator 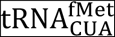efficiently initiates at the UAG amber stop codon from two different reporters, but does not initiate at detectable levels from genomic UAG codons. The amber initiator is thus shown to be orthogonal from a translation initiation point of view. Additionally, we showed that expressed NanoLuc(UAG) reporter proteins contain exclusively methionine at the NY termini, ensuring proteins created from this alternative initiation system have the same properties as wildYtype proteins. We also show for the first time that amber initiator expression causes increased ribosome levels, tRNA degradation, and upregulation of amino acid biosynthetic proteins, although these effects do not seem to affect strain fitness. Together, these results point towards a potential role for the amber initiator tRNA, after additional engineering to reduce its reliance on host aminoacylYsynthetases, in a fully orthogonal translation initiation system.^33^

## METHODS

### Bacterial strains

All designed and constructed plasmids were expressed in the genetically recoded organism *E. coli* strain C321.∆A.*exp*, which was a gift from George Church (Addgene #49018). All strain variants were grown in lysogeny broth Lennox (LB^L^) or on LB^L^ agar, supplemented with zeocin for all *E. coli* strain C321.∆A.*exp* cultures and carbenicillin for cultures with reporter plasmid and spectinomycin for cultures with amber initiator plasmid.

### Plasmid construction

pQEY60::T5Y*sfGFP* reporter plasmids were designed to contain superfolder GFP (sfGFP) with AUG/UAG/GCC start codons. The pULTRA initiator tRNAY expression plasmids were created by using Gibson assembly^34^ to insert a 1971bp gBlock (IDT) contained *metY* genes from MG1655 (GenBank: U00096.3) with anticodon sequences of CAU (wildYtype initiator) and CUA (amber initiator) (Tables S8 and S9) into the pULTRAYCNF^35^ vector (Addgene #48215) (Figure 1A). To create an empty vector, we constructed the pULTRA::*tac*YEmpty plasmid via PCR backbone amplification of pULTRA::*tac2metY*(CUA) and ligation, such that all elements of the pULTRA::*tac2 metY*(CUA) plasmid except the *metY*(CUA) gene were retained. The pSMARTYBAC::*pRha*Y NanoLuc(UAG) and pSMARTYBAC::*pRha*YNanoLuc(AUG) plasmids were constructed by cutting pSMARTYBAC with NotI followed by Gibson assembly with gblocks encoding a rhamnoseYinducible promoter (*pRha*) followed by the NanoLuc genes^36^ with a CY terminal 6xYHis tag and either UAG or AUG start codon. All plasmids were purified and stored in 10 mM TrisYCl, pH 8.5. These plasmids were transformed into C321.ΔA.*exp* strains containing either pULTRA::*tac*Y*metY*(CUA) or pULTRA::*tac*YEmpty.

### Culture growth conditions for assay measurements

Bacterial cultures were grown overnight at 37° C in LB^L^ with appropriate antibiotics. After overnight growth, each culture was diluted 1:100 into 400 µL LB^L^ in a 96Ywell deep well plate containing either 1 mM IPTG to induce, or 2% glucose (w/v) to repress, *metY(CUA)* expression. To control for the effects of the reagents used for repression and induction of the *tacI* promoter, we used the same repressed (2% glucose) and induced (1 mM IPTG) culture conditions for all plasmid combinations, regardless of whether they contained a plasmid with a *tacI*promoter or not.

### Fluorescence measurements

Measurements of fluorescence intensity from the amber initiator plasmid system was performed as before^15^ with the following modifications: OD_600_ was used to estimate culture density, followed by fluorescence measurement (ex=485 nm, em=520 nm, bandwidth = 9 nm, PHERAstar FSX). Cultures from the above experiment were also measured on Cytoflex S flow cytometer (FITC, 488 nm excitation laser, 525/40 nm emission bandYpass filter). 10,000 measured events were triggered on a side scattering threshold. Data was processed using FlowJo v10.

### Fitness analysis

Strains of C321.∆A.*exp* containing the pULTRA::*tac*Y*metY*(CAU), pULTRA::*tac*Y*metY*(CUA), or pULTRA::*tac*YEmpty plasmid were grown overnight as described above. Cell densities were measured, and cells were passaged to a starting OD_600_ of 0.1 into fresh 200 µL LB^L^ with appropriate antibiotics and 1 mM IPTG. Cultures were grown in a flatYbottom 96Ywell plate sealed with gasYpermeable seal at 37°C (300 rpm). Absorbance at OD_600_ over time was measured on a SPECTROstar NANO plate reader at 5Yminute intervals. Growth rate (µ_r_) and maximal cell density (max OD_600_) was determined for each culture using the R package GrowthCurver^37^ represented in Table S2.

### Nanoluciferase Assay

Measurements of luminescence intensity from the amber initiator plasmid system was performed as before^15^ using the NanoYGlo Luciferase Assay System kit (Promega, #N1110) with the following modifications: expression of nanoluciferase was induced with the following concentrations of rhamnose: 0, 5, 50, 100, 500 µM, while 100 μL of culture were mixed with 100 μL of assay buffer and incubated for 15 min at room temperature, followed by luminescence measurement (em=230Y750 nm, gain = 900, PHERAstar FSX).

### NanoLuc expression and purification

Single colonies from C321.ΔA.*exp* strain harbouring pULTRA::*tac*Y*metY*(CUA) and pSMARTYBAC::*pRha*Y*NanoLuc*(TAG) or pSMARTYBAC::*pRha*Y*NanoLuc*(ATG) as a control was used to inoculate 3 mL of LB^L^ supplemented with zeocin, spectinomycin, and chloramphenicol and shaken at 37°C overnight. We used the overnight cultures to purify CYterminal 6X histidineYtagged nanoluciferase protein following NiYNTA purification as before^15^ with the following modifications: expression of 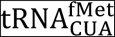and nanoluciferase was induced with 1 mM IPTG and 1 mM rhamnose.

### Sample preparation for mass spectrometry proteome analysis

Strains were grown overnight and back diluted 1:100 into fresh media in 250 mL flasks with appropriate antibiotics and 1 mM IPTG. Cells were harvested three hours and five hours postY induction (3,500 x g/4°C/10 min). Cell pellets were lysed with 0.5 mL CelLytic™ B Cell Lysis buffer (SigmaYAldrich #B7310) adding 0.1 mg lysozyme, 1X protease inhibitor (Roche #04693132001), and 25 units of benzonase (SigmaYAldrich #E1014). The lysate was incubated (15 min/room temperature) then centrifuged (16,000 x g/room temperature/10 min). 1 mL of soluble protein supernatant was collected per culture and analysed with minor changes from previous work.^38^. Proteins from whole cell lysate and NiYNTA purified fractions were precipitated with 1 mL iceYcold acetone (Y 20°C/overnight) then centrifuged (15,000 x g/4°C/10 min), followed by protein pellet resuspension in 100 mM TrisYHCl, 8 M urea (pH 8.0). Protein samples were digested with 1:100 (w/w) protein/trypsin (37°C/overnight), then digested again (37°C/4 hours) with fresh trypsin as before.

### Peptide library generation

To produce a peptide ion library, samples from biological triplicates were pooled and separated by highYpH RPYHPLC as described previously.^39^ Mass spectrometry analysis was performed on a TripleTOF 6600 in dataYdependent mode as previously described^39^ except selection of ion precursors were performed over 140Yminute runs. Spectra were interrogated using ProteinPilot 5.0 using the Paragon algorithm (SCIEX)^17^ and searched against the complete *E. coli* KY12 MG1655 protein sequence database from Uniprot (proteome IDYUP000000625 (last modified 2^nd^ April, 2018)) and inYhouse generated C321.ΔA.*exp* UAG offYtarget database (File S5). The default paragon algorithm includes a fixed cysteine carbamidomethylation modification, caused by dithiothrietol and iodoacetamide treatment, and variable modifications sourced from http://unimod.org.^16^ Proteins and peptides were accepted with a protein false discovery rate of <1%.

### SWATHMMS analysis

Spectra collection was performed in data independent mode as described previously^39^ except, MS/MS spectra were collected across the *m*/*z* range of 350Y1500 with a 35 ms accumulation time. Ion libraries of individual strains were imported into PeakView software 2.1 using the SWATH MicroApp 2.0 (SCIEX) and matched against SWATHYMS data of individual replicates. Retention time calibration was performed using endogenous peptides and data were processed as previously described^39^ to generate cumulative protein areas from extracted ion chromatograms which were exported for further analysis (Figure S3 and S4). Proteins with two or more peptides, pYvalue < 0.01, and foldYchange >2 and <0.5 were considered as significantly differentially expressed (Tables S3 and S4).

### Gene ontology analysis

The differentially expressed proteins were analysed against the total set of 1,803 quantified proteins using the overrepresentation test in PANTHER (http://pantherdb.org) with default parameters.^18^ Gene ontology biological processes with p < 0.05 (after Binomial test with Bonferroni multiple hypothesis test correction) were considered significant.

### Generation of inMhouse offMtarget database

To identify peptides initiating from these genomic UAG codons we generated an offYtarget proteome database using the genome sequence of C321.ΔA.*exp*. We considered all genomic UAG codons as potential start codons and considered UGA and UAA as stop codons. We did not treat UAG as a stop codon because C321.ΔA.*exp* lacks release factor 1 which is required for termination at UAG stop codons. We generated a list of 27,108 ORFs and filtered out all ORFs shorter than 150 nucleotides (50 amino acids) in length. This process resulted in an offYtarget database of 9,857 protein sequences that could potentially form functional ORFs initiating from UAG start codons (Figure 5A). To determine the SDYlike sequences upstream of potential ORFs, we used a previously described method^40^ to predict thermodynamic interactions between the antiYShine Dalgarno and the upstream sequences of the potential ORFs, using the RNA coFold method of the ViennaRNA Package 2.0^41^. We used a similar binding energy threshold of Y4.5 kcal/mol to filter down the offYtarget database, resulting in 990 potential offYtarget proteins with upstream SDYlike sequences (Table S6).

### Targeted protein mass spectrometry of NMterminal peptide and offMtarget proteins

The targeted mass spectrometry method of parallel reaction monitoring (PRM)^31^ was used to identify the NYterminal peptide species of the nanoluciferase reporter protein with a UAG start codon. Additionally, we analysed the whole cell proteome for offYtarget protein expression. Samples were analysed on a high resolution QYExactive mass spectrometer (ThermoFisher Scientific) coupled to an EASYYnLC1000 liquid chromatography system. Peptides were eluted over a 120Yminute linear gradient with increasing concentration of elution buffer (99.9% (v/v) acetonitrile, 0.1% formic acid). The analysis consisted of one survey (full) scan at 70,000 resolution (400 m/z) for targeted precursors (Table S10 and Table S11) (AGC target 2e^5^, maximum injection time 100 ms, 2 m/z isolation width). Measured precursor ions were selected for HCD fragmentation with normalized collision energy of 30 followed by full ms/ms of product ions at 17,500 resolution (m/z 200, AGC target of 2e^5^ and 60 ms maximum injection time). Precursor and fragment ion spectra were analysed using Skyline version 4.2. ^42^ PRM assay result for identification of NYterminal amino acid can be downloaded from the Panorama web repository server (https://panoramaweb.org/H5K69A.url).

## Supporting information

Vincent et al - Amber Initiator - Supporting Information

Vincent et al - Amber Initiator - Supporting Information Tables and Files

## SUPPORTING INFORMATION

Figure S1: Fold change in fluorescence due to 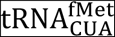induction; Figure S2: Proportion of differentially expressed proteins represented in each gene ontology category; Figure S3: SWATH protein expression profile clusters; Figure S4: Evaluation of SWATH YMS quantification technique; Figure S5: OffYtarget peptide detected in C321.ΔA.*exp* cells regardless of amber initiator expression; Table S1: Translation initiation efficiency; Table S8: Reporter and tRNA gene sequences; Table S9: Synthesized oligonucleotides (PDF).

Table S2: A) Growth curve data represented as OD600 measured every 5 minutes B)Average specific growth rate (min^Y1^) and maximal OD600 (A.U.) (XLSX); Table S3: List of all quantified proteins using SWATHYMS technique three hours postYinduction; Table S4: List of all quantified proteins using SWATHYMS technique five hours postYinduction; Table S5: Gene ontology categorization of differentially expressed proteins five hours postYinduction; Table S6: List of offYtarget ORFs harbouring SDYlike sequences upstream; Table S7: List of all offYtarget proteins harmonized with Uniprot IDs from conventional *E. coli* database; Table S10: Parallel reaction monitoring targeted precursor list for NYterminal amino acid identification; Table S11: Parallel reaction monitoring targeted precursor list for offYtarget peptide identification (XLSX).

Supporting File S1: Amber initiator expression plasmid pULTRA_tacYmetY(CUA) (GB); Supporting File S2: Empty vector control plasmid pULTRA_tacYEmpty (GB); Supporting File S3: GFP reporter plasmid pQEY60_T5YsfGFP(UAG) (GB); Supporting File S4: Nanoluciferase reporter plasmid pSMARTYBAC_pRhaYNanoluc(UAG) (GB); Supporting File S5: FASTA file of combined proteomic database including potential offYtarget proteins for mass spectrometry identification (FASTA).

## ABBREVIATIONS

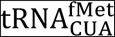: amber initiator tRNA with CUA anticodon

SWATHYMS: Sequential Window Acquisition of all Theoretical fragment ion Mass Spectra

IF1: Initiation Factor 1

IF2: Initiation Factor 2

IF3: Initiation Factor 3

sfGFP(AUG): superfolder green fluorescence protein gene with wildYtype AUG start codon

sfGFP(UAG): superfolder green fluorescence protein gene with amber stop codon UAG as start codon

NanoLuc(AUG): the NanoLuc gene with wildYtype AUG start codon

NanoLuc(UAG): the NanoLuc gene with amber stop codon UAG as start codon

## AUTHOR INFORMATION

### Author Contributions

RMV and PRJ conceived the study. RMV and BWW performed experiments, analyzed data, and created figures and tables. RMV and PRJ drafted the manuscript. All authors read and approved the final manuscript.

## ACKNOWLEDGEMENTS

We thank Matthew McKay, Ardeshir Amirkhani, David Cantor, Karthik Kamath and Dana Pascovici of the Australian Proteome Analysis Facility for assistance in protein and data analysis. We also thank Michael A. Sorensen, Richard Fahlman, Olivier Borkowski, Heinrich Kroukamp, and Jeff Glasgow for helpful discussions. PRJ was supported by the Molecular Sciences Department, Faculty of Science & Engineering, and the Deputy ViceY Chancellor (Research) of Macquarie University. BWW and RMV are recipients of the Macquarie University Research Excellence PhD scholarship (MQRES). Aspects of this research was conducted at the Australian Proteome Analysis Facility, facilitated by the Australian Government’s National Collaborative Research Infrastructure Scheme.

