## Supplementary material for "Measuring amber initiator tRNA orthogonality in a genomically recoded organism": Vincent et al - Amber Initiator - Supporting Information

**CONTENTS**

SUPPORTING FIGURES

Figure S1: Fold change in fluorescence due to $\mathrm{tRNA}^{\mathrm{fMet}}_{\mathrm{CUA}}$ induction.

Figure S2: Proportion of differentially expressed proteins represented in each gene ontology category.

Figure S3: SWATH protein expression profile clusters.

Figure S4: Evaluation of SWATH -MS quantification technique.

Figure S5: Off-target peptide detected in C321.ΔA.*exp* cells regardless of amber initiator expression.

SUPPORTING TABLES

Table S1: Translation initiation efficiency.

Table S2: Average maximal specific growth rate ($\mu_{max}$) and maximal cell density ($OD_{600}$).

Table S3: List of all quantified proteins using SWATH-MS technique three hours post-induction.

Table S4: List of all quantified proteins using SWATH-MS technique five hours post-induction.

Table S5: Gene ontology categorization of differentially expressed proteins five hours post-induction.

Table S6: List of off-target ORFs harbouring SD-like sequences upstream.

Table S7: List of all off-target proteins harmonized with Uniprot IDs from conventional *E. coli* database.

Table S8: Reporter and tRNA gene sequences.

Table S9: Synthesized oligonucleotides.

Table S10: Parallel reaction monitoring targeted precursor list for N-terminal amino acid identification.

Table S11: Parallel reaction monitoring targeted precursor list for off-target peptide identification.

SUPPORTING FILES

Supporting File S1: pULTRA::*tac-metY*(CUA) plasmid.gb

Supporting File S2: pULTRA::*tac*-Empty plasmid.gb

Supporting File S3: pQE-60::T5-*sfGFP*(UAG) reporter plasmid.gb

Supporting File S4: pSMART-BAC::*pRha-NanoLuc*(UAG) plasmid.gb

Supporting File S5: Combined proteomic database including potential off-target proteins.fasta

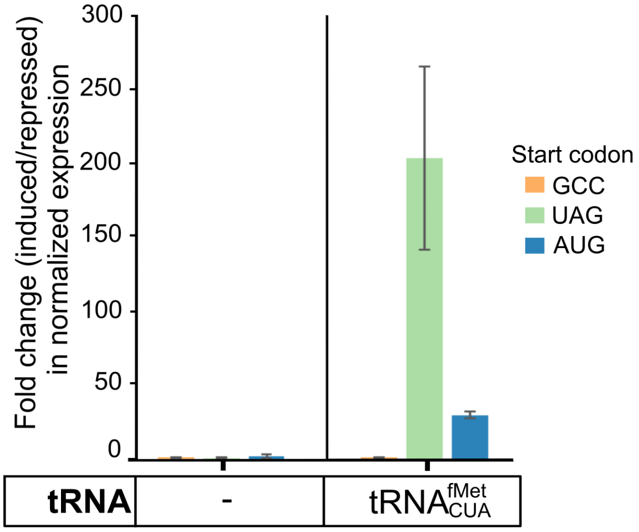


**Figure S1.** Expression of $\mathrm{tRNA}^{\mathrm{fMet}}_{\mathrm{CUA}}$ from pULTRA::*tac-metY*(CUA) amber initiator plasmid causes significantly greater increase in reporter fluorescence from amber UAG start codon versus canonical AUG start codons. Fold change in normalized expression of sfGFP reporter due to IPTG induction. tRNA: presence or absence of $\mathrm{tRNA}^{\mathrm{fMet}}_{\mathrm{CUA}}$ expressing amber initiator plasmid. Average of three biological replicates with error bars showing one standard deviation.

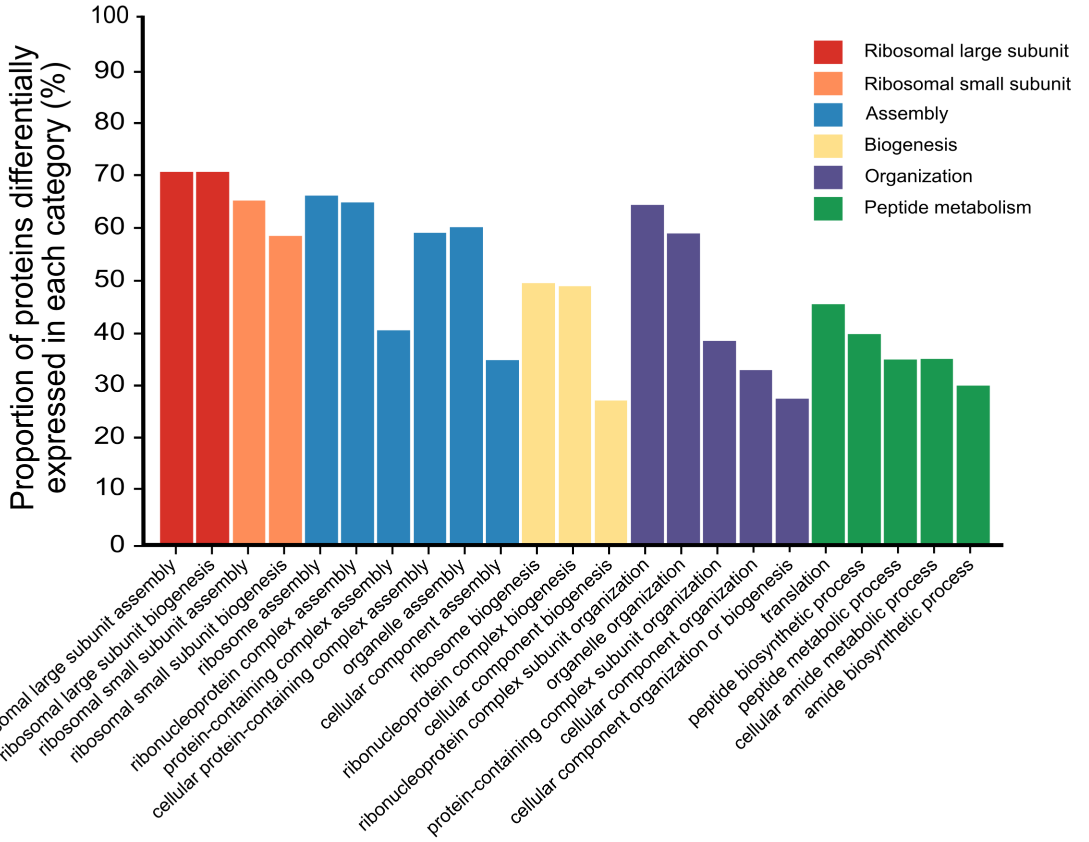


**Figure S2.** Significantly over-represented gene ontology biological process categories for differentially expressed proteins at mid-exponential phase due to $\mathrm{tRNA}^{\mathrm{fMet}}_{\mathrm{CUA}}$ expression. These differentially expressed proteins were compared to 1,803 quantified proteins obtained from SWATH-MS data. The proportion shown is the number of differentially expressed proteins annotated to each gene ontology category divided by the number of quantified proteins linked to at least one annotation term within the indicated gene ontology biological process categories.

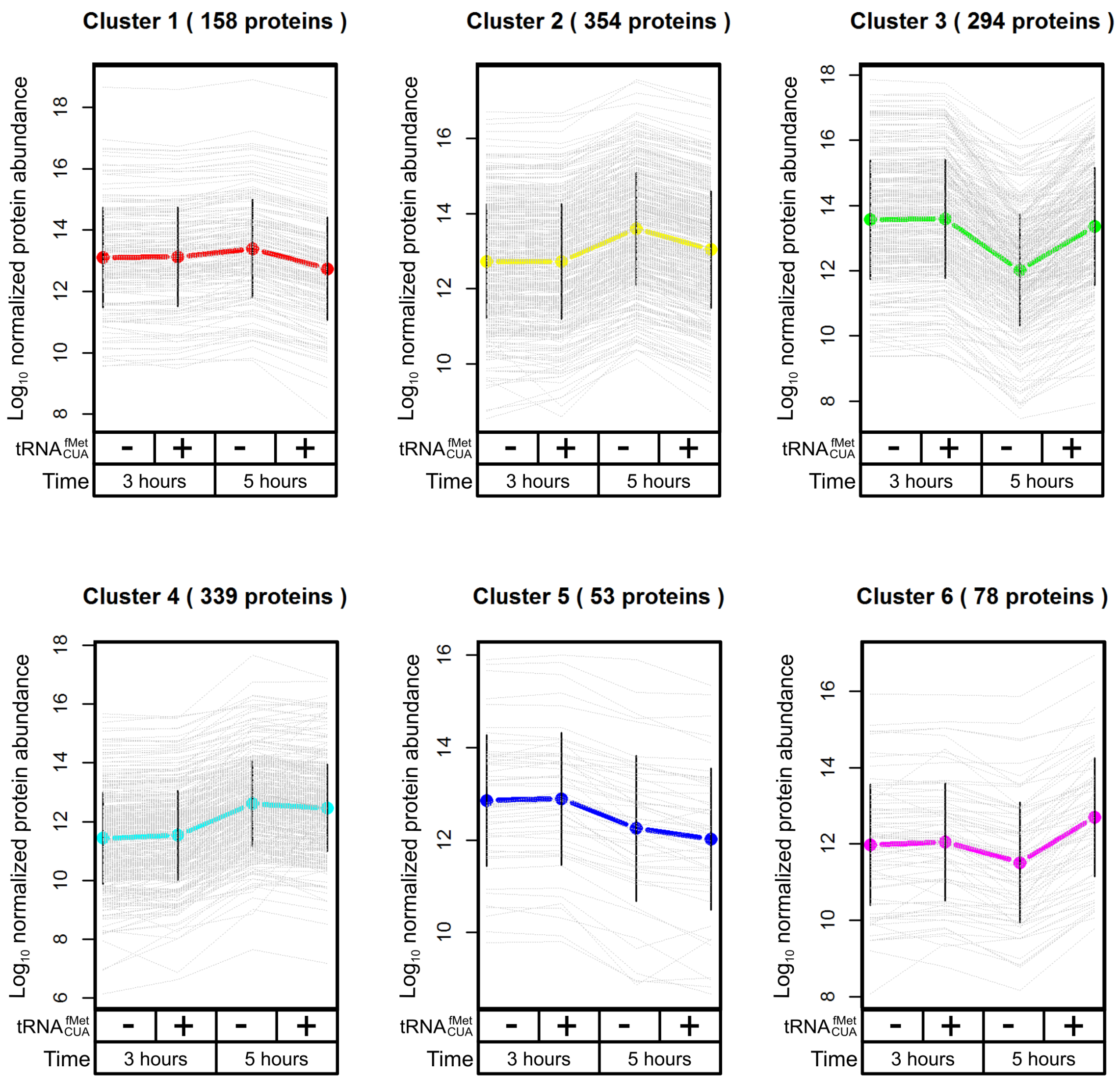


**Figure S3.** Normalized ($log_{10}$) protein expression profile of six clusters of proteins expressing or lacking $tRNA^{fMet}_{CUA}$ at three and five hours post-induction with 1 mM IPTG. A total of 1,276 proteins were divided into six clusters according to their expression profile.

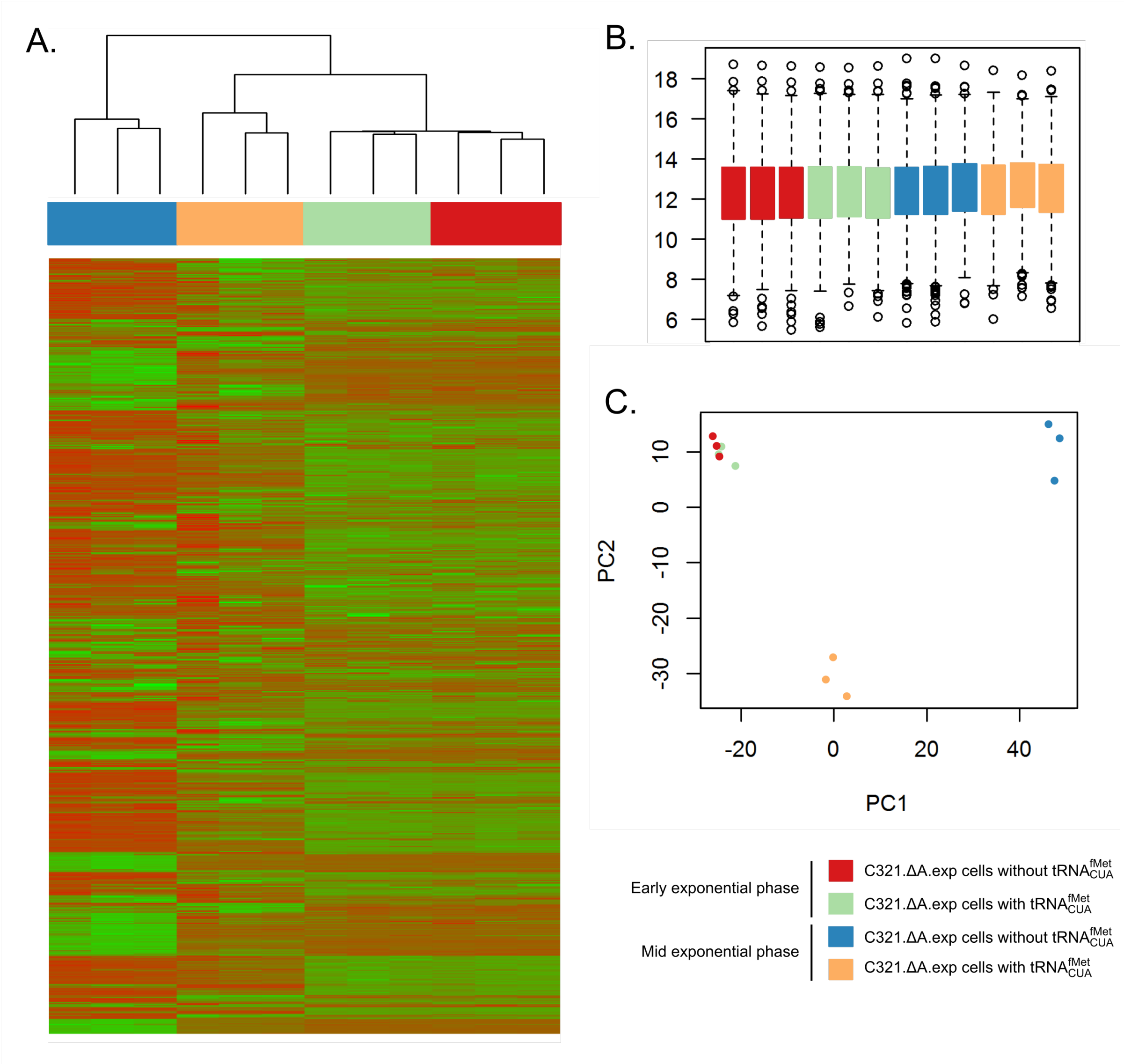


**Figure S4.** Evaluation of SWATH-MS quantification technique. (A) The hierarchical clustering heat map of SWATH-MS log transformed protein expression data of C321.ΔA.*exp* cells with or without tRNA$^{fMet}_{CUA}$ at early exponential phase and mid exponential phase. (B) Box plots of log transformed normalized protein peak areas of individual biological replicates displaying reproducibility from SWATH-MS. (C) Strains of C321.ΔA.*exp* with and without tRNA$^{fMet}_{CUA}$ at different growth phase. Samples classified based on log transformed and normalized SWATH-MS protein expression values using Principal component analysis (PCA). Each point represents a biological replicate.

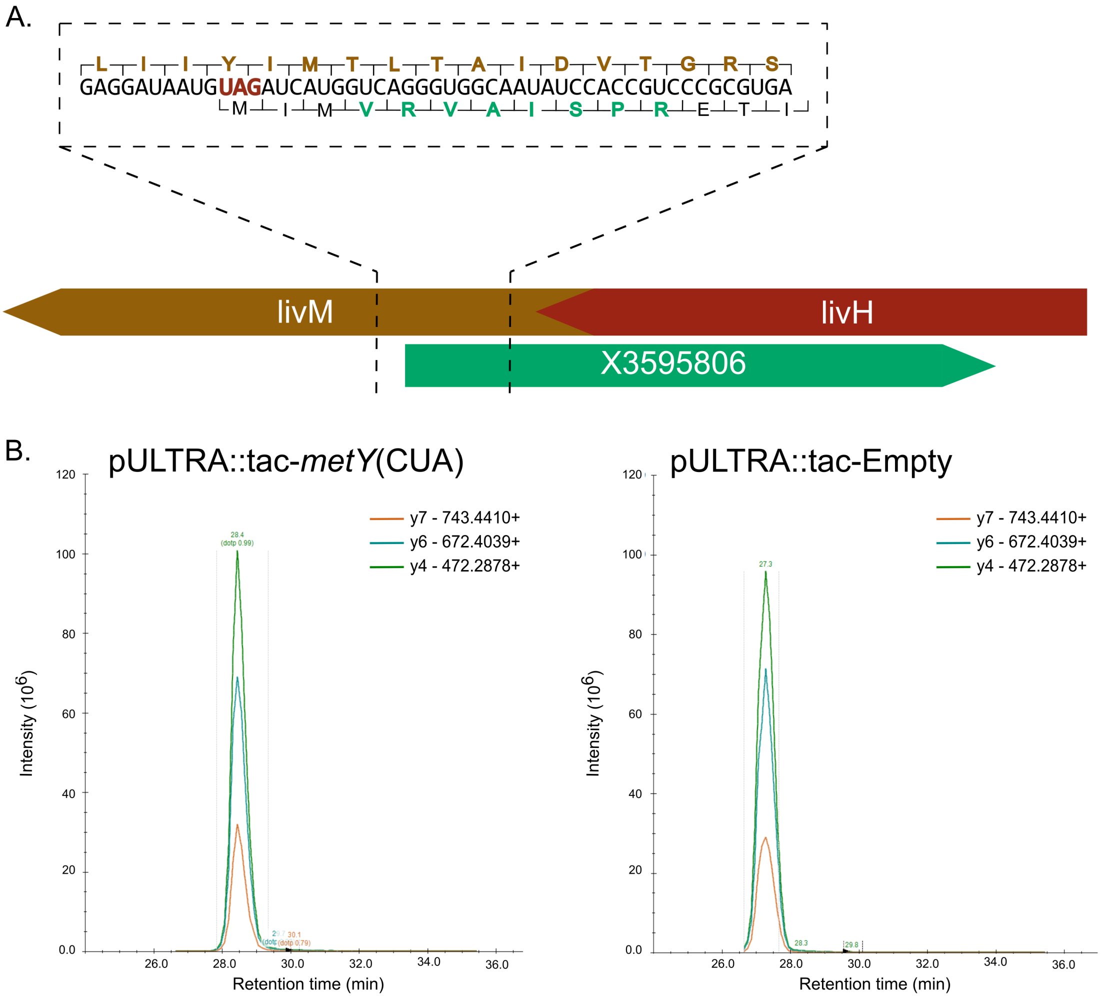


**Figure S5.** Off-target peptide detected in C321.ΔA.*exp* cells regardless of amber initiator expression. (A) Schematic diagram representing off-target ORF X3595806 placement with respect to *E. coli* host genes *livM* and *livH*. The dotted box depicts the host *livM* gene sequence in black, partial host protein in brown, the partial off-target protein sequence in black, and the peptide observed via PRM in green. (B) Peptide from off-target protein X3595806 is expressed in empty vector control. Product ions of the peptide observed in the presence of amber initiator plasmid (left) or the empty vector (right).

**Table S1.** Translation initiation efficiency (%). Average normalized fluorescence (Arb. U./$OD_{600}$) represented in Figure 1B was calculated from three biological replicates. Translation initiation efficiency was determined by comparing the average normalized fluorescence of cultures expressing sfGFP with individual start codons to the average normalized fluorescence of cultures without the amber initiator plasmid expressing sfGFP with an AUG start codon, in respective growth conditions.

| **Amber initiator plasmid** | **Growth condition** | **Start codon** | **Average normalized fluorescence (Arb. U./$OD_{600}$)** | **Standard deviation** | **Translation initiation efficiency (%)** |
|---|---|---|---|---|---|
| No plasmid | Repressed | AUG | 5319.12 | 2119.0 | 100.0 |
| | | UAG | 49.35 | 7.7 | 0.9 |
| | | GCC | 41.15 | 1.9 | 0.8 |
| pULTRA::*tac-metY*(CUA) | Repressed | AUG | 3738.70 | 580.4 | 70.3 |
| | | UAG | 51.48 | 9.7 | 1.0 |
| | | GCC | 44.27 | 0.9 | 1.2 |
| No plasmid | Induced | AUG | 11793.56 | 4329.3 | 100.0 |
| | | UAG | 58.20 | 13.7 | 0.5 |
| | | GCC | 62.68 | 13.9 | 0.5 |
| pULTRA::*tac-metY*(CUA) | Induced | AUG | 112895.55 | 7786.1 | 957.3 |
| | | UAG | 10503.45 | 2518.1 | 89.1 |
| | | GCC | 66.40 | 39.9 | 0.6 |

**Table S8: Reporter and tRNA gene sequences.**

| Gene | Sequence |
|---|---|
| $\mathrm{tRNA}^{\mathrm{fMet}}_{\mathrm{CUA}}$ | CGCGGGGTGGAGCAGCCTGGTAGCTCGTCGGGCTCTAAACCCGAAGATCGTCGGTTCAAA<br>TCCGGCCCCCGCAACCA |
| sfGFP(AUG) | ATGCGTAAAGGCGAAGAGCTGTTCACTGGTGTCGTCCCTATTCTGGTGGAACTGGATGGT<br>GATGTCAACGGTCATAAGTTTTCCGTGCGTGGCGAGGGTGAAGGTGACGCAACTAATGGT<br>AAACTGACGCTGAAGTTCATCTGTACTACTGGTAAACTGCCGGTACCTTGGCCGACTCTGG<br>TAACGACGCTGACTTATGGTGTTCAGTGCTTTGCTCGTTATCCGGACCATATGAAGCAGCA<br>TGACTTCTTCAAGTCCGCCATGCCGGAAGGCTATGTGCAGGAACGCACGATTTCCTTTAAG<br>GATGACGGCACGTACAAAACGCGTGCGGAAGTGAAATTTGAAGGCGATACCCTGGTAAA<br>CCGCATTGAGCTGAAAGGCATTGACTTTAAAGAAGACGGCAATATCCTGGGCCATAAGCT<br>GGAATACAATTTTAACAGCCACAATGTTTACATCACCGCCGATAAACAAAAAAATGGCATT<br>AAAGCGAATTTTAAAATTCGCCACAACGTGGAGGATGGCAGCGTGCAGCTGGCTGATCAC<br>TACCAGCAAAACACTCCAATCGGTGATGGTCCTGTTCTGCTGCCAGACAATCACTATCTGA<br>GCACGCAAAGCGTTCTGTCTAAAGATCCGAACGAGAAACGCGATCATATGGTTCTGCTGG<br>AGTTCGTAACCGCAGCGGGCATCACGCATGGTATGGATGAACTGTACAAA |
| sfGFP(UAG) | TAGCGTAAAGGCGAAGAGCTGTTCACTGGTGTCGTCCCTATTCTGGTGGAACTGGATGGT<br>GATGTCAACGGTCATAAGTTTTCCGTGCGTGGCGAGGGTGAAGGTGACGCAACTAATGGT<br>AAACTGACGCTGAAGTTCATCTGTACTACTGGTAAACTGCCGGTACCTTGGCCGACTCTGG<br>TAACGACGCTGACTTATGGTGTTCAGTGCTTTGCTCGTTATCCGGACCATATGAAGCAGCA<br>TGACTTCTTCAAGTCCGCCATGCCGGAAGGCTATGTGCAGGAACGCACGATTTCCTTTAAG<br>GATGACGGCACGTACAAAACGCGTGCGGAAGTGAAATTTGAAGGCGATACCCTGGTAAA<br>CCGCATTGAGCTGAAAGGCATTGACTTTAAAGAAGACGGCAATATCCTGGGCCATAAGCT<br>GGAATACAATTTTAACAGCCACAATGTTTACATCACCGCCGATAAACAAAAAAATGGCATT<br>AAAGCGAATTTTAAAATTCGCCACAACGTGGAGGATGGCAGCGTGCAGCTGGCTGATCAC<br>TACCAGCAAAACACTCCAATCGGTGATGGTCCTGTTCTGCTGCCAGACAATCACTATCTGA<br>GCACGCAAAGCGTTCTGTCTAAAGATCCGAACGAGAAACGCGATCATATGGTTCTGCTGG<br>AGTTCGTAACCGCAGCGGGCATCACGCATGGTATGGATGAACTGTACAAA |
| sfGFP(GCC) | GCCCGTAAAGGCGAAGAGCTGTTCACTGGTGTCGTCCCTATTCTGGTGGAACTGGATGGT<br>GATGTCAACGGTCATAAGTTTTCCGTGCGTGGCGAGGGTGAAGGTGACGCAACTAATGGT<br>AAACTGACGCTGAAGTTCATCTGTACTACTGGTAAACTGCCGGTACCTTGGCCGACTCTGG<br>TAACGACGCTGACTTATGGTGTTCAGTGCTTTGCTCGTTATCCGGACCATATGAAGCAGCA<br>TGACTTCTTCAAGTCCGCCATGCCGGAAGGCTATGTGCAGGAACGCACGATTTCCTTTAAG<br>GATGACGGCACGTACAAAACGCGTGCGGAAGTGAAATTTGAAGGCGATACCCTGGTAAA<br>CCGCATTGAGCTGAAAGGCATTGACTTTAAAGAAGACGGCAATATCCTGGGCCATAAGCT<br>GGAATACAATTTTAACAGCCACAATGTTTACATCACCGCCGATAAACAAAAAAATGGCATT<br>AAAGCGAATTTTAAAATTCGCCACAACGTGGAGGATGGCAGCGTGCAGCTGGCTGATCAC<br>TACCAGCAAAACACTCCAATCGGTGATGGTCCTGTTCTGCTGCCAGACAATCACTATCTGA<br>GCACGCAAAGCGTTCTGTCTAAAGATCCGAACGAGAAACGCGATCATATGGTTCTGCTGG<br>AGTTCGTAACCGCAGCGGGCATCACGCATGGTATGGATGAACTGTACAAA |

**Table S9: Synthesized oligonucleotides.**

| **Gibson assembly oligonucleotides** | |
|---|---|
| 548-pULTRA_back-FOR | CCTGTCAGTAACGAGCAGCAATAGACATAAGCGGCTATTTAACGACCC |
| 549-pULTRA_back-REV | GGAAAGATGAACGTGATGATGCCCTTGAGAGCCTTCAACCC |
| 550-*metY*(CUA)_2-FOR | GGCTCTCAAGGGCATCATCACGTTCATCTTTCCCTGGTTGCC |
| 551-*metY*(CUA)_2-REV | CCGCTTATGTCTATTGCTGCTCGTTACTGACAGGAAAATGGGCAGCC |
| **Screening and sequencing oligonucleotides** | |
| 552-Amber_screen-FOR^ | TGAACCGCTCTAGATTTCAGTG |
| 553-Amber_screen-REV^ | AGATCCGGCCACGATGAC |

**^PCR with these primers gave a 0.7 kb product for pULTRA::tac-*metY*(CUA).**
